# Opioid receptor architecture for the modulation of brainstem functions

**DOI:** 10.1101/2022.12.24.521865

**Authors:** Nicholas F. Hug, Nicole Mercer Lindsay, William M. McCallum, Justin Bryan, Karen Huang, Nicole Ochandarena, Adrien Tassou, Grégory Scherrer

## Abstract

Opioids produce profound and diverse effects on a range of behaviors, many driven by brainstem activity; however, the presence of opioid and opioid-like receptors at this level has been poorly studied outside of nociceptive structures and components of respiratory circuitry. While previous studies identified expression of µ, δ, κ, and nociceptin opioid and opioid-like receptors in the brainstem, patterns have not been fully delineated, and neither has receptor coexpression nor the behavioral implications of their expression in most structures. We aimed to elucidate expression patterns for all four receptors across somatosensory-motor, auditory, and respiratory brainstem circuits; identify recurring themes to provide insight into the mechanisms by which exogenous opioids affect broader brainstem circuits; and characterize the function of endogenous opioids in subcortical processing and behavior modulation. Using a fluorescent reporter mouse line for each receptor, we created a comprehensive atlas of brainstem receptor distribution and identified novel expression patterns in modality-specific circuits. Each receptor showed unique expression patterns across the brainstem with minimal correlation between receptors. Orofacial somatosensory-motor circuits expressed all four receptors, though generally in distinct regions, suggesting differential opiate modulation of afferent and efferent trigeminal signaling. Within the auditory circuits, receptors segregated along the vertical and horizontal processing pathways with minimal colocalization. Finally, the respiratory circuit strongly expressed the µ opioid receptor in multiple crucial structures with minimal presence of the other three receptors. We further assessed the functional significance of these expression patterns, using the respiratory circuitry as an example, by characterizing respiratory responses to selective opioid agonists, finding that each agonist caused unique alterations in breathing pattern and/or breath shape. Together, these results establish a comprehensive atlas of opioid and opioid-like receptor expression throughout the brainstem, laying the essential groundwork for further evaluation of opioid neuromodulation across the broad spectrum of behaviors.

## Introduction

Opioids are an invaluable component of pain treatment in a wide variety of clinical settings. For example, opioid medications are critical for the induction and maintenance of anesthesia, controlling postoperative pain, reducing recovery time and hospital stay duration, relieving the severe pain associated with chronic conditions such as cancer, and facilitating comfort during end-of-life care. When used appropriately and carefully monitored, opioids provide safe and reliable pain management. However, their potential for abuse, addictive properties, and overdose, have led to a concerning rate of non-medical or illicit consumption of opioids and contributed to an immense number of deaths over the past few decades in the United States. From 1999 to 2011, consumption of both prescription and non-prescription opioids increased by as much as 500% (Kolodny et al., 2015). In 2010 alone, prescription opioids contributed to over 16,000 deaths in the U.S. (Dart et al., 2015). Furthermore, the center for disease control and prevention (CDC) calculations of yearly opioid overdoses increased by approximately 20% in the U.S. between 2019 and 2020 (Ahmad et al. 2021). As this class of medications is essential and unrivaled in their ability to manage pain, preventing their significant adverse events is crucial for continued safe use. Accomplishing this requires elucidation of the mechanisms underlying opioid function and side effects.

While studies emphasize the analgesic and addictive properties of opioids, their diverse effects (e.g., nausea, constipation, tolerance, hyperalgesia, respiratory depression) reflect the broad expression of opioid and opioid-like receptors (ORs) in the central, peripheral, and enteric nervous system (Pert and Snyder, 1973; Bagnol et al., 1997; Basbaum et al., 2009; Erbs et al., 2015; Mosińska et al., 2016; Wang et al., 2018). Exogenous and endogenous opioids act on four distinct Class A G protein-coupled receptors (GPCRs): the three canonical μ, δ, and κ opioid receptors (MOR, DOR, KOR) and the structurally related but pharmacologically distinct nociceptin receptor (NOP) (Corder et al., 2018). Activation of any of these receptors can result in G_i_ protein signaling and activation of arrestin-dependent pathways (Manglik et al., 2016). Despite their structural and signaling similarity, activation of each of the four receptors results in diverse downstream effects in response to opioid ligands. Previous studies reported OR expression in an array of brain structures, including throughout the brainstem (Erbs et al., 2015), suggesting endogenous and exogenous opioids influence a broader range of behaviors than those currently under exploration.

The presence of ORs in the brainstem has received little attention beyond structures involved in nociception (e.g., the parabrachial nucleus (PB), periaqueductal gray (PAG), rostral ventromedial medulla (RVM)), and components of the respiratory circuitry (Bachmutsky et al., 2020; Varga et al., 2020; Liu et al., 2021). Previous work revealed the coexpression of MOR and DOR in more than 50 brainstem structures, suggesting that these two receptors cooperate (Erbs et al., 2015). KOR and NOP expression has also been studied in the brainstem to a lesser degree (Mansour et al., 1994; Anton et al., 1996). Importantly, the coexpression of all four receptors has not been investigated, nor has the behavioral or clinical relevance of OR expression in most brainstem regions. Thus, we aimed to elucidate expression patterns for MOR, DOR, KOR, and NOP across somatosensory-motor, auditory, and respiratory brainstem circuits; identify recurring themes to provide insight into the mechanisms by which exogenous opioids affect broader brainstem circuits; and characterize the function of endogenous opioids in subcortical processing and behavior modulation.

To accomplish these goals, we resolved brainstem expression of all four receptors using a fluorescent reporter mouse line for each receptor (Scherrer et al., 2006; Erbs et al., 2015; Mann et al., 2019; Chen et al., 2020). First, we created a three-dimensional (3D) atlas of receptor distribution in the brainstem. Second, we identified novel receptor expression patterns within the somatosensory-motor, auditory, and respiratory circuits. Finally, we determined the behavioral implications of the differential expression patterns observed in these circuits and demonstrated their functional significance within the respiratory circuit. This comprehensive atlas of receptor expression throughout the brainstem lays the essential groundwork for functional studies for the opioid neuromodulation of vital brainstem functions.

## Results

### General principles of opioid receptor expression

We first sought to characterize OR expression patterns throughout all brainstem circuits. We used *Oprm1*^*mCherry/mCherry*^ (MORmCherry) (Erbs et al., 2015), *Oprd1*^*GFP/GFP*^ (DORGFP) (Scherrer et al., 2006), *Oprk1*^*tdTomato/tdTomato*^ (KORtdTomato) (Chen et al., 2020), and *Oprl1*^*YFP-floxYFP-flox*^ (NOPYFP) (Mann et al., 2019) mice to visualize each of the four receptors. We quantified the distribution of each fluorescently labeled receptor across 80 brainstem nuclei, 4 cerebellar regions, and, to contextualize brainstem expression, 9 forebrain regions (Berg et al., 2019; Groeneboom et al., 2020; Yaoyao, 2020) (**Fig 1A, B**, see Methods). We observed three overarching themes for brainstem expression: 1) regionally restricted expression of MOR and DOR, 2) sparse expression of KOR, and 3) broad NOP expression across structures (**Fig 1B,C**).

**Figure 1.**
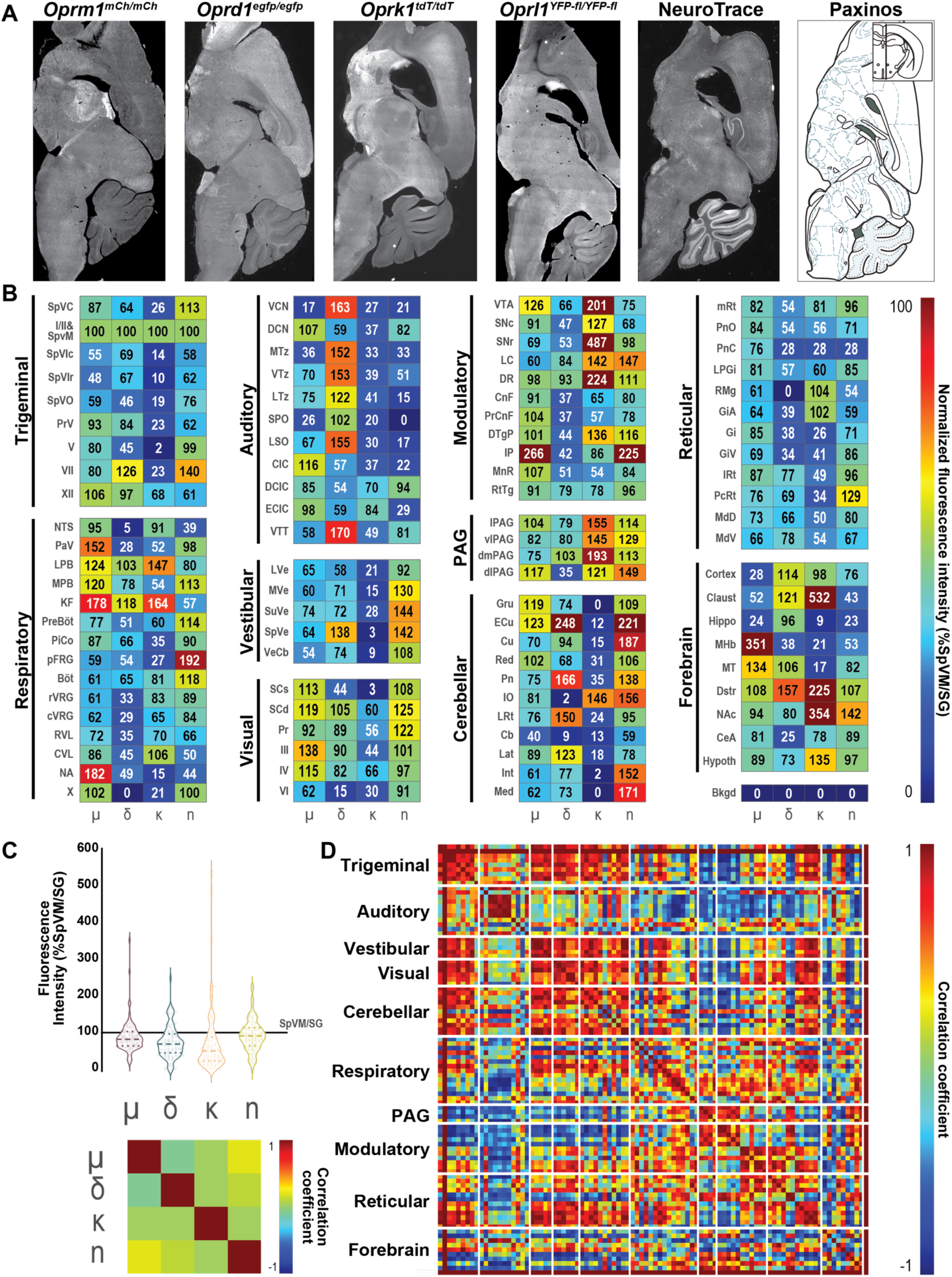
Opioid receptor expression profiles across brainstem circuits. **(A)** Example sagittal images showing, left to right: *Oprm1*^*mCherry/mCherry*^ (MORmCherry), *Oprd1*^*egfp/egfp*^ (DORGFP), *Oprk1*^*tdT/tdT*^ (KORtdTomato), *Oprl1*^*YFP-fl/YFP-fl*^ (NOPYFP), NeuroTrace counterstain, and a Paxinos atlas section. **(B)** Quantification of fluorescence intensity of 81 brainstem, 4 cerebellar, and 9 forebrain structures for MOR (μ), DOR (δ), KOR (κ), and NOP (n), normalized between background (0) and fluorescence intensity in the medullary dorsal horn lamina I/II and SpVM (100) across the 94 structures measured. **(C, top)** Violin plots of fluorescence intensity for all measured structures shows the distribution of intensity for each OR. Pairwise correlations between ORs **(C, bottom)** and brain regions **(D)**.

We first show unique expression patterns of each OR in the brainstem (**Fig 1C,D**). We found that the brain structures with the highest fluorescence intensity for MOR and KOR include previously studied forebrain regions, the medial habenula (MHb) and claustrum (Claust), respectively. However, MOR is also expressed abundantly across brainstem nuclei, notably in the paratrigeminal (PaV), lateral parabrachial (LPB), Kölliker-Fuse (KF), and nucleus ambiguus (NA), regions controlling respiratory, cardiac, and other vagal functions (Ramirez et al., 2021). In contrast, KOR expression varies widely, with some brain regions expressing KOR at very high intensity compared to most others. These regions include the ventral tegmental area (VTA), substantia nigra pars reticulata (SNr), and dorsal raphe (DR) (**Fig 1B,C**). We also observe statistically significant KOR expression in 78% of brainstem regions (p < 0.05, **Fig S1**), with particularly notable KOR intensity in the LPB, KF, and PAG (**Fig 1B**), which suggests KOR may influence brainstem signaling in addition to its known functions in motivation and reward.

In contrast to MOR and KOR, DOR and NOP have their highest fluorescence intensity in brainstem regions, the external cuneate nucleus (ECu) and interpeduncular nucleus (IP), respectively. We also observe strong DORGFP fluorescence in auditory nuclei (e.g., ventral cochlear nucleus (VCN), medial nucleus of the trapezoid body (MTz)), brainstem regions that project to the cerebellum (the pontine nuclei (Pn) and ECu), and in the lateral deep cerebellar nucleus (Lat) (**Fig 1B**). While NOP has relatively consistent bright fluorescence intensity across the brainstem, we observe notable NOP expression in the vestibular nuclei, dorsolateral PAG (dlPAG), ECu, and cuneate nucleus (Cu) (**Fig 1B,C**).

Each receptor displays a unique brain-wide expression pattern, with the only significant correlation being observed between MOR and NOP (including all quantified regions, r = 0.20, p < 0.05) (**Fig 1C**). Surprisingly, few brain regions show statistically significant correlation patterns with other regions (**Figs 1D, S2**). The rarity of receptor pattern correlation between brain regions demonstrates that expression profiles from all four receptors rarely repeat in the exact same proportions (**Fig 1D,E**). However, related regions sometimes display similarities, typically involving one or two of the four receptors. For example, receptor expression patterns throughout the spinal trigeminal nuclei (i.e., secondary sensory nuclei for somatosensation and nociception for the face) highly correlate (R>0.71 for each pair). These nuclei include the spinal trigeminal nucleus pars caudalis (SpVC), muralis (SpVM), interpolaris caudalis (SpVIc), interpolaris rostralis (SpVIr), and oralis (SpVO) (**Figs 1D, S2**, p < 0.05). Similarly, receptor expression significantly correlates among brainstem regions involved in vestibular information processing, including between secondary or tertiary structures, as well as with the Int and Med deep cerebellar nuclei (**Figs 1D, S2**, p < 0.05). In contrast, OR expression profiles for regions involved in motivation and reward tend to highly express KOR (e.g., NAc or SNr) and anticorrelate with other brain regions (**Figs 1D, S2**).

### Trigeminal somatosensory-motor processing

Based on our observations regarding OR expression correlations between trigeminal nuclei (**Fig 1C,E**), we further clarified expression patterns of orofacial somatosensory-motor circuits. Trigeminal complex nuclei and cranial motor nuclei together mediate low-level sensory processing and execution of reflexes and simple movements (Mercer Lindsay et al., 2019; Ruder et al., 2021). In contrast to auditory and visual stimuli, somatosensation primarily conveys information detailing the environment of the animal’s peri-personal space (McElvain et al., 2018). Orofacial somatosensation often utilizes fine movements of the sensors themselves (e.g., tongue licking or vibrissa whisking), which necessitates a close association between sensory and motor circuit components (Moore et al., 2015; McElvain et al., 2018). While previous work has extensively studied dorsal horn circuits, much less is known about the opioid receptor expression patterns throughout the trigeminal nuclei and their functional implications.

Sensory neurons innervating the facial skin, gums, teeth, and orofacial mucosa (i.e., nasal, oral, and lingual mucosa) transmit signals to the trigeminal nuclei that line the lateral edge of the pons and medulla (**Fig 2A**). The types and densities of somatosensory neuron endings in the skin (e.g., Merkel cells, lanceolate endings, free nerve endings) have specialized arrangements serving unique functions in different facial tissues; notable examples include the innervation of the eye, tongue, and rodent vibrissa. Regardless of this diversity, all facial Aβ somatosensory neurons send information through parallel collateral branches into multiple subdivisions of the trigeminal nuclei, including a population that collateralizes to PrV, SpVO, SpVIr, SpVIc, and SpVC (**Fig 2A**), albeit with distinct bouton patterns in each (Jacquin et al., 1993; Miyoshi et al., 1994; Furuta et al., 2006). In turn, each trigeminal subnucleus projects to an array of unique targets and serves diverse functional roles. First, SpVC and SpVM (sometimes called the SpVI/SpVC transition zone) receive nearly all nociceptive, TrpV1+ afferents for the face (Bae et al., 2004), in addition to low threshold mechanoreceptor afferents (LTMRs), for orofacial pain perception (Chattipakorn et al., 1999; Nash et al., 2009; Shimizu et al., 2009; Castro et al., 2017). SpVC contains a similar layering pattern to that of the dorsal horn and is often called the medullary or trigeminal dorsal horn (Miyoshi et al., 1994; Pradier et al., 2019). Second, PrV and SpVIc neurons receive afferent input from non-overlapping Aβ fibers which they in turn transmit to ventral posterior thalamus (Bosman et al., 2011; Moore et al., 2015). The small receptive fields of PrV and SpVIc neurons result in highly segregated somatotopic maps that are established in these brainstem nuclei and continue up to the VP thalamus and primary somatosensory cortex (e.g., the barrel cortex). Last, SpVIr and SpVO receive afferents from Aβ trigeminal ganglion neurons that in turn project to motoneurons and are associated with orofacial reflexive movements (Morcuende et al., 2002; Takemura et al., 2006; Bellavance et al., 2017).

**Figure 2.**
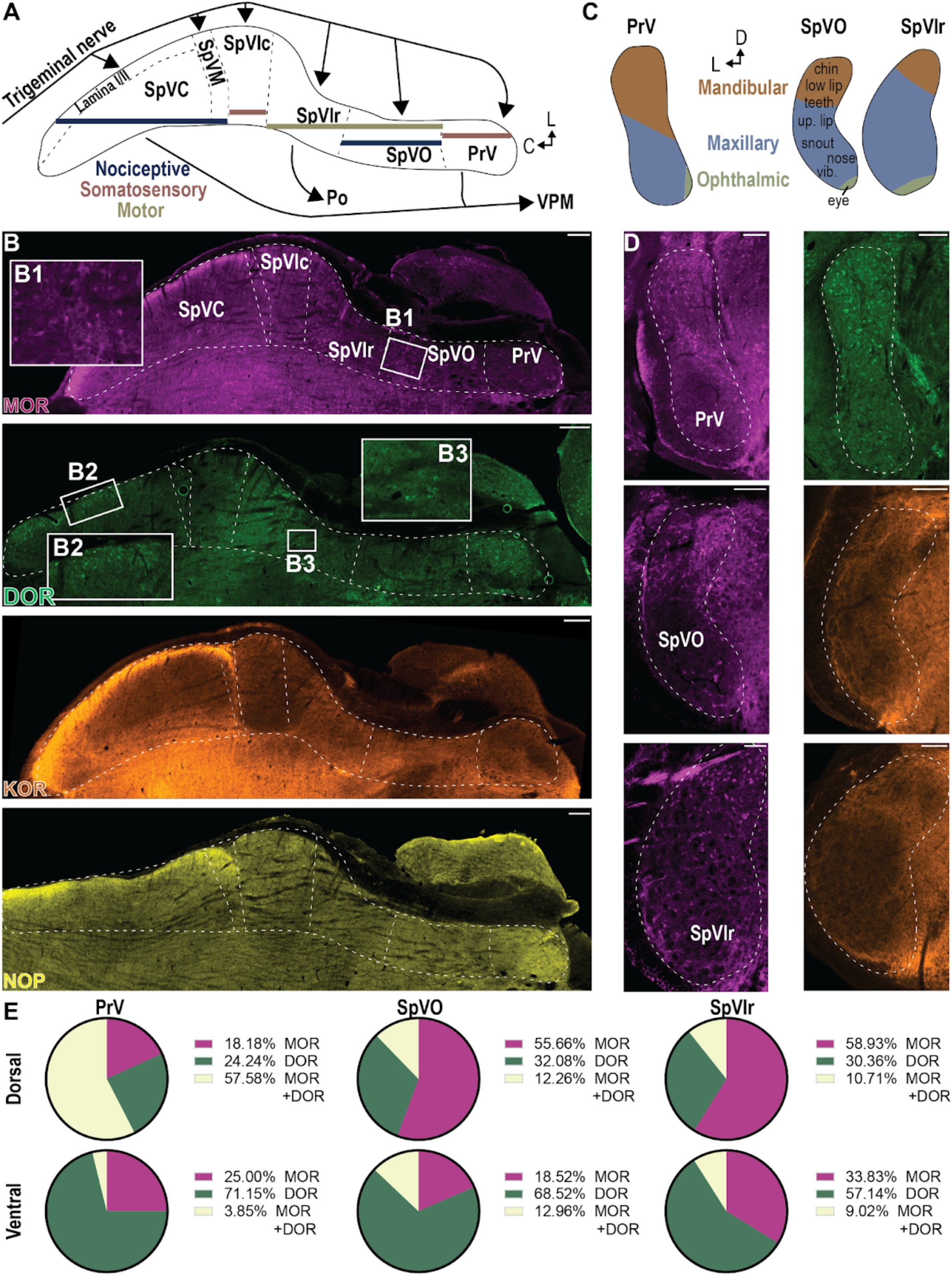
Expression of opioid receptors in trigeminal nuclei varies in relation to subnucleus identity and somatotopic map. **(A)** Schematic of a horizontal section of the trigeminal nuclei SpVC, SpVM, SpVIc, SpVIr, SpVO, PrV. **(B)** Example horizontal sections showing MOR, DOR, KOR, and NOP expression across the trigeminal nuclei in (A). **(C)** Schematic representation of the distribution of mandibular, maxillary, and ophthalmic afferent inputs into PrV, SpVO, and SpVIr. **(D)** Coronal images show OR expression in PrV, SpVO, and SpVIr across the mandibular, maxillary, and ophthalmic subdivisions. **(E)** Pie charts quantifying MOR+, DOR+, and MOR+/DOR+ neuronal populations in PrV, SpVO, and SpVIr.

We observe MOR, DOR, KOR, and NOP in lamina I/II of SpVC (**Fig 2B**), similar to observations in the superficial dorsal horn (Wang et al., 2018). Further, SpVM, a region with particularly dense innervation of the oral and nasal mucosa as well as the cornea, shows strong expression of MOR, KOR, and NOP (**Fig S3A**). Thus, these receptors may mediate somatosensory processing for the mucosa. We observe MOR, KOR, and NOP labeling in fibers in both SpVC and SpVM and, in addition, MOR and DOR labeling in cell bodies (**Figs 2B, S3B**). The trigeminal ganglia, like dorsal root ganglia, contain cells that collectively express all four receptors; these labeled fibers likely correspond to these sensory inputs.

Trigeminal nerve afferent inputs to PrV, SpVO, SpVIr, and SpVIc (**Fig 2A**) maintain a relatively similar somatotopic organization (**Fig 2C**). The dorsal, ventral, and central aspects of these nuclei process mandibular (e.g., jaw, chin, lower lip, and lower teeth), ophthalmic (e.g., eye, cornea, periocular skin), and maxillary somatosensation (e.g., upper lip, snout, nose, and vibrissa), respectively (**Fig 2C**) (Bosman et al., 2011; Matthews, 2012; Matthews et al., 2015). Across the entire somatotopic representation and throughout these four subdivisions, we observe DOR+ neurons (**Fig 2B,D,G**). Additionally, we find that, in contrast to DOR, expression of MOR and KOR is restricted to the dorsal mandibular and ventral ophthalmic subregions of PrV, SpVO, and SpVIr (**Fig 2B-F**). MOR+/DOR+ neurons are rare (between 3 and 12% of the combined MOR+ and DOR+ populations) except in dorsal PrV (mandibular representation), where over 50% of MOR+ and over 50% DOR+ neurons coexpress the other receptor (**Fig 2G**).

Together, these data suggest that, for the primary trigeminal thalamic mechanosensory signaling pathways (e.g., PrV to ventral posteromedial (VPM) thalamus), MOR and DOR modulate afferent signals from the jaw, tongue, and lower lip. However, DOR alone acts on inputs from the rest of the face (**Fig 2G**). Close examination of SpVO and SpVIr suggests that MOR and KOR could modulate premotor, reflexive, and local brainstem circuits for the jaw, tongue, lower lips, and eyelids (e.g., blinking) (**Fig 2E,F**).

Cranial nerve motoneurons innervate the jaw, nose, vibrissa, ear, eyelid, and tongue. Three cranial motor nuclei give rise to these motoneurons: the trigeminal, facial, and hypoglossal nuclei (V, VII, and XII, respectively) (**Fig 3A**). Motoneurons for each muscle cluster together, forming myotopic maps (**Fig 3B-C**). To label motoneurons for both intrinsic and extrinsic muscles, we injected CTB into the vibrissa pad, where the muscles interweave to facilitate protraction and retraction of the vibrissa (int and ex in **Fib 3B**). We use the distribution of CTB neurons, along with previous work, to identify locations of motoneurons for the nasolabialis profundus nose muscle (Deschênes et al., 2016), the orbicularis oculi eyelid muscle (Morcuende et al., 2002; Sun, 2012), the buccinator and levator labii lip muscles (Furutani et al., 2004), and the anterior and posterior auricular levators, and transverse auricular ear muscles (Friauf and Herbert, 1985) (nlp, oo, buc, lev, mla, mlp, and mta, respectively, in **Fig 3B**). While we observed no OR fluorescence in the digastric or masseter divisions of V (**Fig S4**), we did identify motoneurons expressing either MOR or DOR throughout VII, with MOR found primarily in the medial subregions (e.g., the lip and ear motoneurons) and DOR more laterally (e.g., the vibrissa and nose motoneurons), a segregated pattern of expression that mimics the distinct segregation of DOR and MOR in ventral horn motoneurons (Wang et al. 2018). KOR+ fibers intersperse between cell bodies in VII, between the lip and extrinsic vibrissa (retractor) motoneurons (**Fig 3D**).

**Figure 3.**
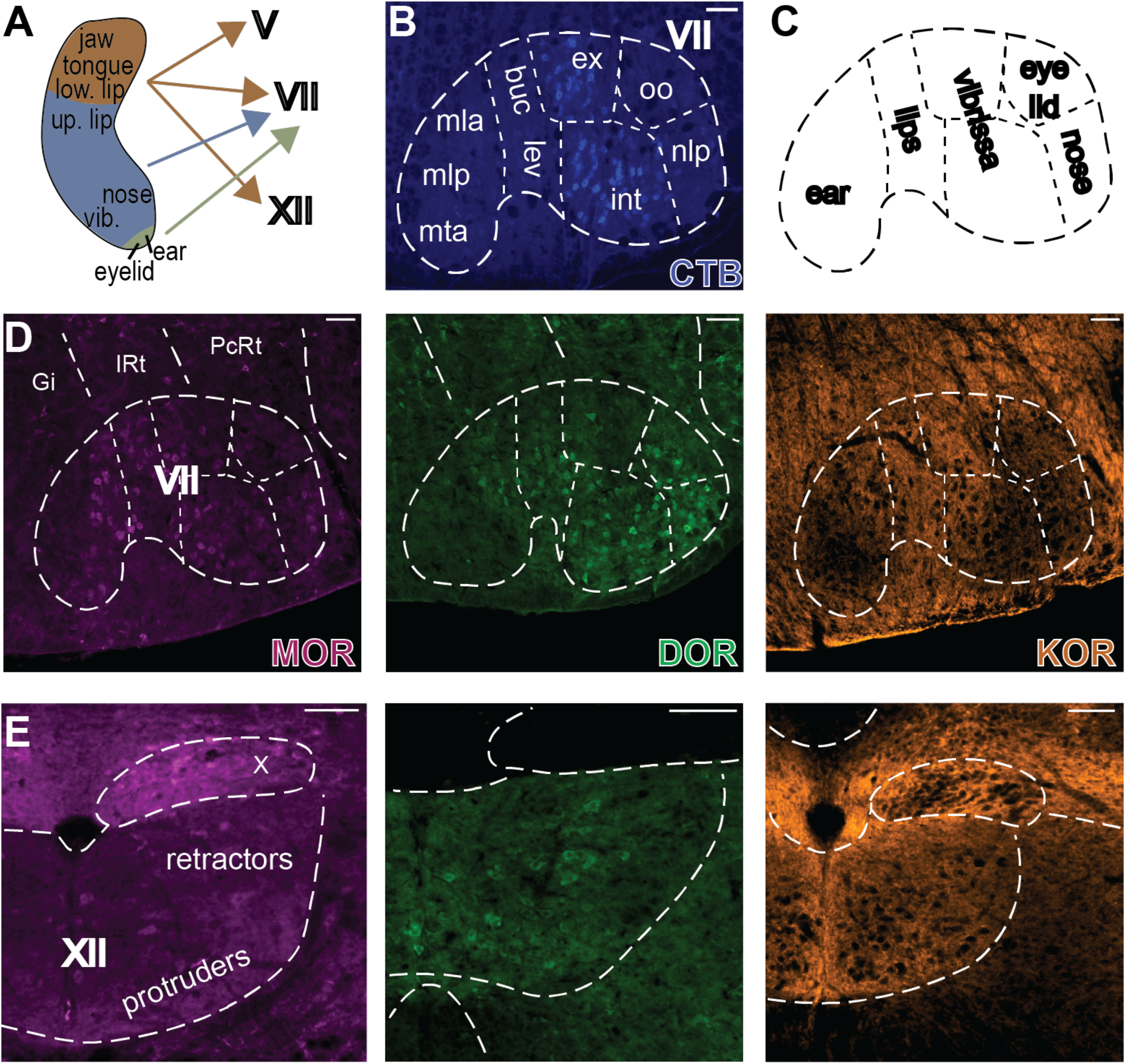
MOR, DOR, and KOR expression in cranial motoneurons. **(A)** Premotor neuron distribution in SpV. **(B)** CTB retrograde tracing from the vibrissa pad to align histological sections with the known myotopic map **(C)** in cranial motor nucleus VII. Nasolabialis profundus (nlp) motoneuron distribution identifies the location of the nose (Deschênes et al. 2016); orbicularis oculi (oo) for the eye (Morcuende et al. 2002); intrinsic and extrinsic for the vibrissa (int and ex) <u>(Takatoh et al. 2013; Takatoh et al. 2021)</u>; buccinator and levator labii for the lip (buc and lev) (Furutani et al. 2004); and anterior/ posterior auricular levators, and transverse auricular for the ear (mla, mlp, and mta) (Friauf and Herbert 1985). **(D)** MOR, DOR, and KOR distribution in VII. Parvocellular reticular formation (PcRt), Intermediate reticular formation (IRt), Gigantocellular reticular formation (Gi). **(E)** MOR, DOR, and KOR distribution in XII. Scale bars indicate 50 µm.

XII innervates the tongue muscles, consisting of tongue protruders and tongue retractors. Ventral XII contains motoneurons for the protruders, while retractors originate in the dorsal XII (**Fig 3E**) (Wang et al., 2020). MOR and DOR scatter sparsely among XII motoneurons (**Fig 3E**). The ventral XII also displays KOR+ fibers (**Fig 3E**). The tongue contracts rhythmically with the respiratory cycle; however, while previous research described the impact of MOR in respiratory circuits, the potential effects of OR agonists on the tongue and resulting changes in respiration remains unclear. Given our findings, we propose that MOR, DOR, and KOR signaling may reduce tongue posture or movement. In concert with tongue inhibition, these data suggest that MOR and KOR activity may also suppress or refine ear, tongue, and vibrissa movements. DOR is positioned to modulate tongue, lip, vibrissa, and nose posture or movements. Exogenous opioids impact locomotion in rodents; however, here we present data suggesting that these same opioid agonists may also influence orofacial motor actions.

### μ and δ opioid receptors segregate to vertical or horizontal auditory circuits

The auditory brainstem receives input from the cochlea, initiates sound localization and coincidence detection computations, and mediates low-level auditory-motor functions. Both illicit and prescription opioid use can induce temporary or permanent hearing damage (Yorgason et al., 2010; Lopez et al., 2012; Hughes et al., 2022). While the mechanism underlying opioid-induced hearing loss remains unclear, cochlear implants that directly activate the nerve can successfully restore function, suggesting an involvement of the cochlea and early brainstem processing (**Fig 4A**).

**Figure 4.**
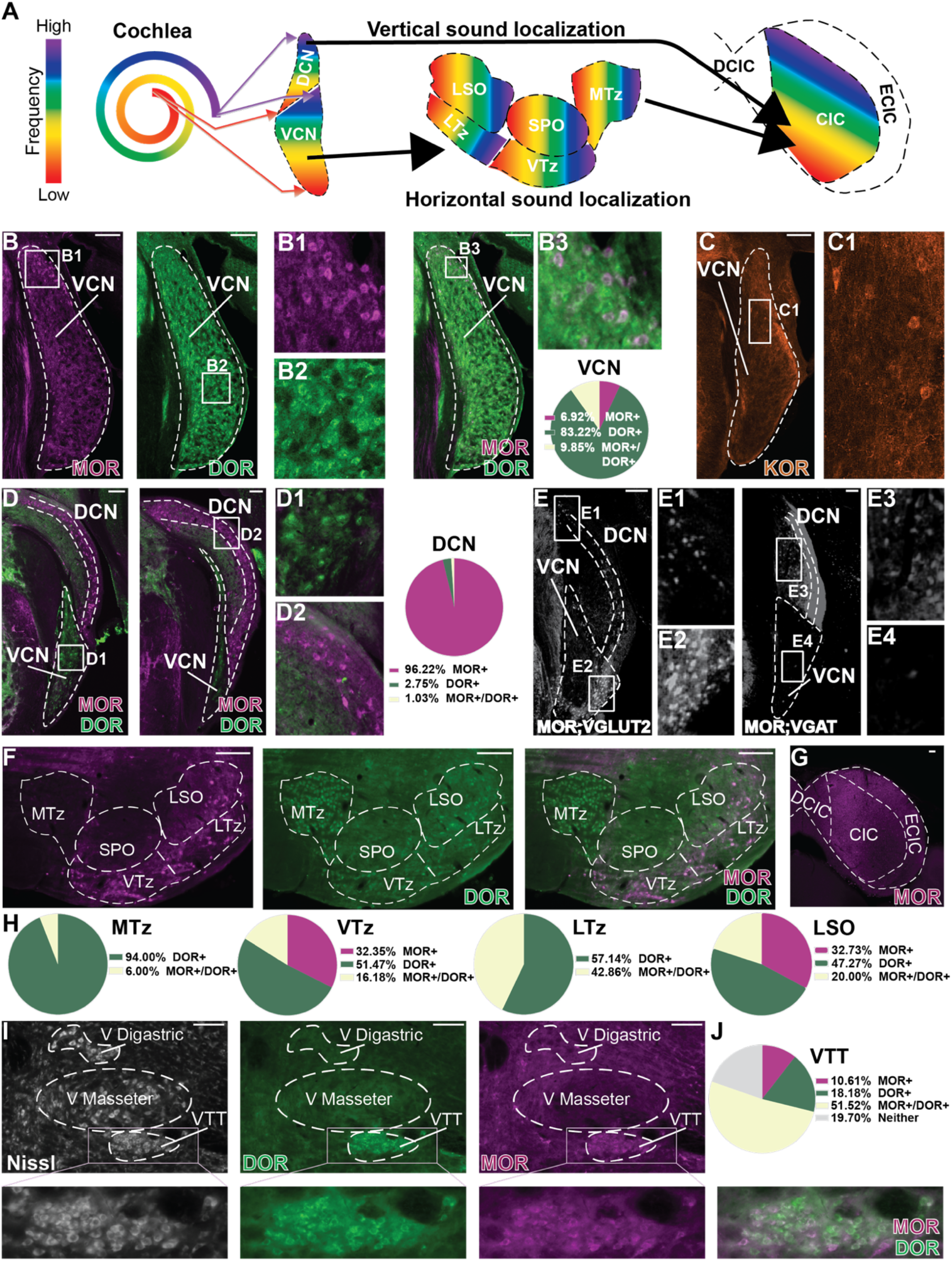
MOR and DOR segregate to vertical and horizontal sound localization circuits in the brainstem. **(A)** Schematic of brainstem auditory circuits, including the orientation of the tonotopic map. **(B)** MOR and DOR distribution in VCN highlighting coexpression in the granule cell layer which, at this rostro-caudal location (approximately Bregma −5.40mm), is located dorsally. **(C)** KOR+ cell bodies in the granule cell layer of VCN. **(D)** MOR and DOR expression in DCN and the caudal VCN (approximately Bregma ∼6.12mm caudal). **(E)** Triple-transgenic mice MOR-Cre; VGLUT2-Flp; Ai65D and MOR-Cre; VGAT-Flp; Ai65D label MOR+ excitatory and inhibitory neurons, respectively, in the DCN and VCN. **(F)** Superior olivary and periolivary nuclei predominantly express DOR with some MOR. **(G)** MOR expression in IC. **(H)** Quantification of MOR and DOR overlap in (F). **(I)** MOR and DOR expression patterns in cranial motor nucleus V. **(J)** Quantification of MOR and DOR expression in the tensor tympani motoneurons (VTT). Scale bars indicate 50 µm.

The cochlear nucleus receives tonotopically matched input from the cochlea. It comprises a laminated, cerebellum-related dorsal nucleus (DCN), including a mix of excitatory and inhibitory neurons, as well as a primarily excitatory ventral nucleus (VCN) with a rind-and-core organization (Kandler et al., 2009; Kandel et al., 2021). DCN and VCN receive highly organized cochlear inputs, with afferent fibers making precise connections to specific cell types in DCN or VCN (Oliver et al., 2018). These nuclei perform initial computations to determine horizontal and vertical sound localization (i.e., whether a sound comes from one’s right or left or above or below them) and coincidence detection (assessing the temporal association of sounds arriving at different ears). They then convey this information to the superior olivary complex (SOC), inferior colliculus (IC), and superior colliculus (SC) in parallel (Kandel et al., 2021). These pathways together are critical for identifying sounds, their source, and appropriate head orientation.

We found that DCN and VCN display contrasting OR expression patterns, with VCN expressing abundant DOR and DCN predominantly expressing MOR (**Fig 4B,D**). DOR+ VCN neurons are scattered throughout the anterior-posterior distribution (**Fig 4D1**) with a slight decrease visible in the posterior part, called the octopus cell area, which mediates coincidence detection and expresses DOR to a lesser degree (**Fig 4D1**). In contrast, MOR+ VCN neurons are restricted almost exclusively to the granule cell layer (**Fig 4B1,B3**). The vast majority of VCN neurons, including granule cells, express either VGLUT1 or VGLUT2 (Oliver and Cant, 2018), indicating that MOR and DOR here are likely expressed in glutamatergic neurons. In addition to MOR+ and MOR+/DOR+ neurons, the granule cell layer of VCN further includes KOR+ neurons (**Fig 4C**) with the potential for triple coexpression of MOR, DOR, and KOR. Granule cell layer neurons primarily project from VCN to DCN (Oliver et al., 2018), positioning ORs to presynaptically modulate neurotransmission between the VCN and DCN at this early processing level.

In contrast to VCN, DCN contains abundant MOR+ neurons, but minimal DOR+ neurons. In particular, we observe MOR+ neurons across the fusiform layer, the primary computational layer of DCN that projects to the CIC (**Fig 4D**). The two halves of the cochlear nucleus mediate different aspects of hearing: While VCN calculates horizontal sound localization and coincidence detection, DCN computes vertical sound localization. The clearly segregated expression of DOR and MOR to the VCN and DCN, respectively, positions each receptor to influence separate dimensions of sound localization.

To determine if MOR+ neurons in the DCN and VCN granule cell layer are excitatory or inhibitory, we used a Cre- and Flp-dependent mouse reporter line crossed with *Oprm1*^*Cre/+*^ and either *Slc17a6*^*FLPo/+*^ (MOR-Cre;VGLUT2-FLPo) or *Slc32a1*^*FLPo/+*^ (MOR-Cre;VGAT-FLPo) (**Fig 4E**). Surprisingly, we observed almost no MOR+/VGLUT2+ neurons in the DCN (**Fig 4E,E1**) and instead saw, in the MOR; VGAT mice, recapitulation of the fluorescence pattern observed in MORmCherry mice (**Fig 4E,E3**). Using these same mice, we confirmed that almost all VCN granule cells expressing MOR are also VGLUT2+ (**Fig 4E,E2,E4**).

The superior paraolivary nucleus (SPO) and periolivary nuclei receive input from the VCN (**Fig 4A**) and then compute horizontal sound origin via two methods, depending on sound frequency: 1) for low-frequency sounds, by identifying the difference in latency of a sound to arrive at each cochlea; or 2) for high-frequency sounds, by comparing sound intensity between the two cochleae and VCNs (Kandel et al., 2021). VCN inputs comprise a web of unilateral, contralateral, or bilateral connections to specific SPO and/or periolivary nuclei that compare miniscule latency differences (Oliver et al., 2018). Notably, the medial, ventral, and lateral trapezoid nuclei (MTz, VTz, LTz) are primarily glycinergic and play an important role in rapidly inhibiting the circuit (Oliver and Cant, 2018). In mice, MTz is the largest of these nuclei and is densely DOR+ with scant expression of MOR (**Fig 4F,H**). In contrast, the smaller VTz and LTz lie ventral to SPO, medial superior olive (MSO), and lateral superior olive (LSO), and sparsely express DOR and MOR (**Fig 4F,H**). The OR expression pattern of LSO appears similar to that of VTz and LTz, with occasional MOR+/DOR+ neurons (**Fig 4F,H**). Thus, MOR and DOR may cooperate in VTz, LTz, and LSO. However, the dense presence of DOR+ neurons in MTz, the primary inhibitor of the SOC in mice, more strongly implies that activation of DOR may impede horizontal sound localization. Interestingly, this mirrors the OR pattern at the level of the cochlear nuclei, where, as discussed above, DOR+ neurons dominate VCN, which is crucial for horizontal sound processing.

Vertical sound localization circuits originate in DCN and project directly to the central nucleus of IC (CIC) (**Fig 4A**). The external cortex of the IC (ECIC) integrates inputs primarily from somatosensory brainstem regions and conveys these multimodal signals to CIC (Beyerl, 1978; Oliver and Cant, 2018; Kandel et al., 2021). IC, broadly, integrates all auditory signals from brainstem nuclei before subsequent transfer to the medial geniculate thalamic nucleus for thalamocortical auditory processing. Notably, only MOR is strongly present in CIC, yet restricted to the medial half, suggesting a role for MOR in modulating auditory processing across the entire tonotopic map (Kandel et al., 2021) (**Fig 1F**). This expression pattern mimics findings at the level of the cochlear nuclei in which MOR+ neurons predominate at the site of vertical sound localization, the DCN.

Finally, the inner ear has two muscles, the tensor tympani and the stapedius, innervated by efferent motoneurons originating in the trigeminal motor nucleus (VTT) and facial motor nucleus (VIIst), respectively.

These muscles facilitate a reflexive response to intense acoustic stimuli by increasing impedance in the middle ear, protecting against hearing loss (Mukerji et al., 2010). VTT motoneurons distinctly express MOR and DOR, with a high percentage of overlap (**Fig 4I,J**), suggesting that MOR and DOR agonists may suppress the intensity of sounds by acting directly on inner ear motoneurons.

Overall, we find notable MOR and DOR expression across brainstem auditory regions in a pattern correlating with distinct functions and, further, that MOR and DOR rarely colocalize within auditory regions. KOR and NOP are expressed more sparsely in the auditory regions; however, when present, they colocalize with other ORs (**Fig S1**). Importantly, we observe the second instance of an organization that we have found to repeat throughout the brainstem: secondary sensory nuclei, here VCN and DCN, typically express one dominant OR. A circuit-level perspective reveals a second pattern specific to the auditory system: DOR expression in VCN and regions of SOC that mediate horizontal sound localization, versus MOR expression in DCN and CIC, the pathway responsible for vertical sound localization (Oertel and Fay, 2013). However, coexpression of MOR and DOR is common in VTz, LTz, and VTT motoneurons. Altogether, this evidence suggests that DOR and MOR both modulate sound localization, but predominantly in different spatial dimensions. While there are occasional instances of coexpression, we expect the overall behavioral outcome to be consistent, with DOR and MOR playing discrete functions in sound modulation.

### Respiratory and other vagal circuits strongly express μ opioid receptor and lowly express the δ, κ, and nociceptin opioid receptors

Though studies exploring opioid-induced respiratory depression (OIRD) have investigated components of the brainstem circuits responsible for respiratory patterning, opioid receptor expression throughout the circuit lacks detailed characterization. This complex circuit consists of dorsal, ventral, and pontine respiratory groups with highly complex interactions between these groups as well as with cortical and peripheral involvement.

The dorsal respiratory group (DRG) comprises the nucleus tractus solitarius (NTS), paratrigeminal nucleus (PaV), and secondary sensory nuclei. These regions receive inputs from the vagus nerve, which transmits nociceptive, pulmonary, cardiac, gastrointestinal, and gustatory signals (**Fig 5A**). The NTS viscerotopic map (i.e., topographic pattern of visceral afferent input) evolves rostro-caudally, with the rostral region receiving gustatory information, followed by gastrointestinal, then cardiac, and pulmonary most caudally (Mazzone and Undem, 2016; Cutsforth-Gregory and Benarroch, 2017). The PaV map also evolves rostro-caudally with nociceptive vagal inputs most rostrally followed by overlapping cardiac and pulmonary inputs (Caous et al., n.d.) (**Fig 5A**). (Caous et al., n.d.; Cutsforth-Gregory and Benarroch, 2017)

**Figure 5.**
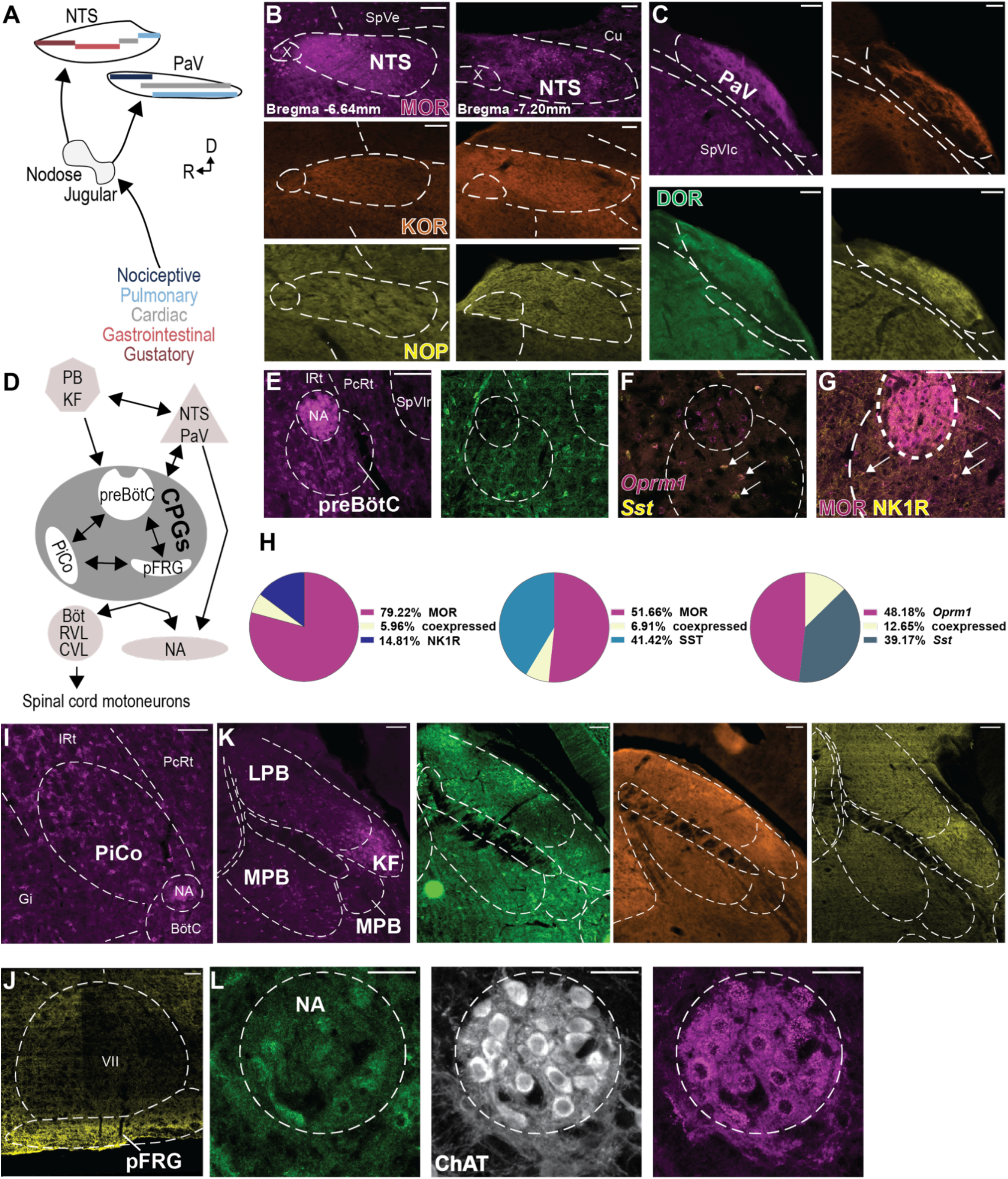
Respiratory circuits predominantly express MOR, with limited DOR, KOR, and NOP expression. **(A)** Schematic of vagus afferent input patterns into NTS and PaV through the nodose and jugular ganglia. **(B)** NTS expression of MOR, KOR, and NOP in a rostral (Bregma −6.64mm) and caudal (Bregma −7.2mm) section. **(C)** PaV expression of all four ORs. **(D)** Schematic of respiratory circuits including the dorsal respiratory group (DRG includes NTS and PaV), ventral respiratory group (VRG includes the CPGs (preBӧtC, pFRG, and PiCo), BӧtC, RVL, CVL, and NA), and the pontine respiratory group (PRG includes PB and KF). **(E)** Expression of MOR and DOR in preBӧtC. **(F)** *In situ* hybridization (ISH) showing coexpression of MOR RNA (*Oprm1*) and somatostatin RNA (*Sst*) in preBӧtC. **(G)** MOR and NK1R coexpression in preBӧtC. **(H)** Quantification of MOR coexpression with NK1R, SST (antibody), and *Sst* (ISH). **(I)** MOR expression in PiCo. **(J)** NOP expression in pFRG. **(L)** MOR and DOR expression in NA. Scale bars indicate 100µm.

Secondary sensory neurons in NTS and PaV project across the brainstem, with inputs to the other two members of the respiratory triad: the ventral respiratory group (VRG), including the central pattern generators (CPGs) for respiratory rhythm, and pontine respiratory group (**Fig 5H**). The ventral respiratory group primarily controls respiratory rhythm generation via three distinct CPGs: the preBötzinger complex (preBötC), parafacial respiratory group (pFRG), and post-inspiratory complex (PiCo) (Del Negro et al., 2018) (**Fig 5H**). preBötC, a small, heterogeneous nucleus found ventral to NA and caudal to the Bötzinger complex (BötC), drives the critical inspiratory breathing pattern. Secondary to preBötC, PiCo neurons drive the post-inspiratory phase of respiration (Anderson et al., 2016; Toor et al., 2019) and pFRG, located ventral to VII, generates the rhythm of active expiration (Del Negro et al., 2018). We find that subpopulations of preBötC and PiCo neurons express MOR, with negligible expression of other ORs (**Fig 5D,E**). The pFRG, in contrast, expresses only NOP (**Fig 5F**). Although the number of neurons expressing MOR in preBötC appears relatively low, selective removal of MOR from this brainstem region partially ameliorates OIRD (Bachmutsky et al., 2020; Varga et al., 2020), suggesting that MOR+ neurons impact the function of this primary respiratory CPG. NOP activation in pFRG could alter pattering of active expiration; however, activity of this CPG only drives rhythm secondary to preBötC’s inspiratory rhythm, suggesting a minimal effect on respiratory patterning compared to MOR activation.

Beyond the three CPGs, the VRG also includes rhythm-supporting nuclei like BӧtC and airway premotor neurons in the rostral and caudal ventral respiratory groups (rVRG and cVRG), retrotrapezoid nucleus (RTN), and NA. Collectively, these nuclei regulate all features of the respiratory rhythm (Del Negro et al., 2018). BötC modulates the inspiratory-expiratory phase transition, rVRG contains inspiratory premotor neurons, and cVRG provides a premotor drive for expiration (Del Negro et al., 2018). BötC, rVRG, and cVRG all contain MOR+ neurons, albeit somewhat sparsely (**Fig S6**).

The third part of the respiratory triad, the pontine respiratory group (PRG) comprising PB and KF, provides top-down regulation of the transition between inspiration and expiration (Dutschmann and Dick, 2012; Varga et al., 2020). Lateral PB (LPB) densely expresses all four ORs, with particularly robust DOR expression dorsally, and elevated MOR expression in the external LPB subdivision (**Fig 5G**) (Wang et al., 2018), suggesting that MOR and DOR act on complimentary but distinct circuits. KF, found immediately ventrolateral to LPB, expresses all four receptors more strongly than LPB (**Fig 5G**). The medial PB (MPB) expresses MOR, KOR, and NOP but very little DOR. Like those in the preBötC, knockout studies in these nuclei have shown partial respiratory recovery following morphine administration (Varga et al., 2020).

Given the differential distribution of the receptors in respiratory brainstem regions, we aimed to provide an example of the translation implications of our histological analysis by evaluating how their activation impacts behavior. FIrst, we measured the baseline respiratory rate of freely moving wild-type mice using whole-body plethysmography for thirty minutes (**Fig 6A**) and then for sixty minutes after systemic injection of either saline or selective MOR, DOR, KOR, or NOP agonists (20 mg/kg morphine, 5 mg/kg SNC80, 6 mg/kg (±)-U-50488, or 10 mg/kg SCN221510) (**Fig 6B,C**). Consistent with the literature, we observed robust OIRD after morphine administration (**Fig 6D-F**), with wider breath shapes (**Fig 6G**), increased breath duration (**Fig 6H**), and decreased breath amplitude (**Fig 6I**) compared with vehicle-administered mice. These data indicate that MOR activation slows the respiratory rhythm by increasing the duration of each breath, especially the inspiratory phase.

**Figure 6.**
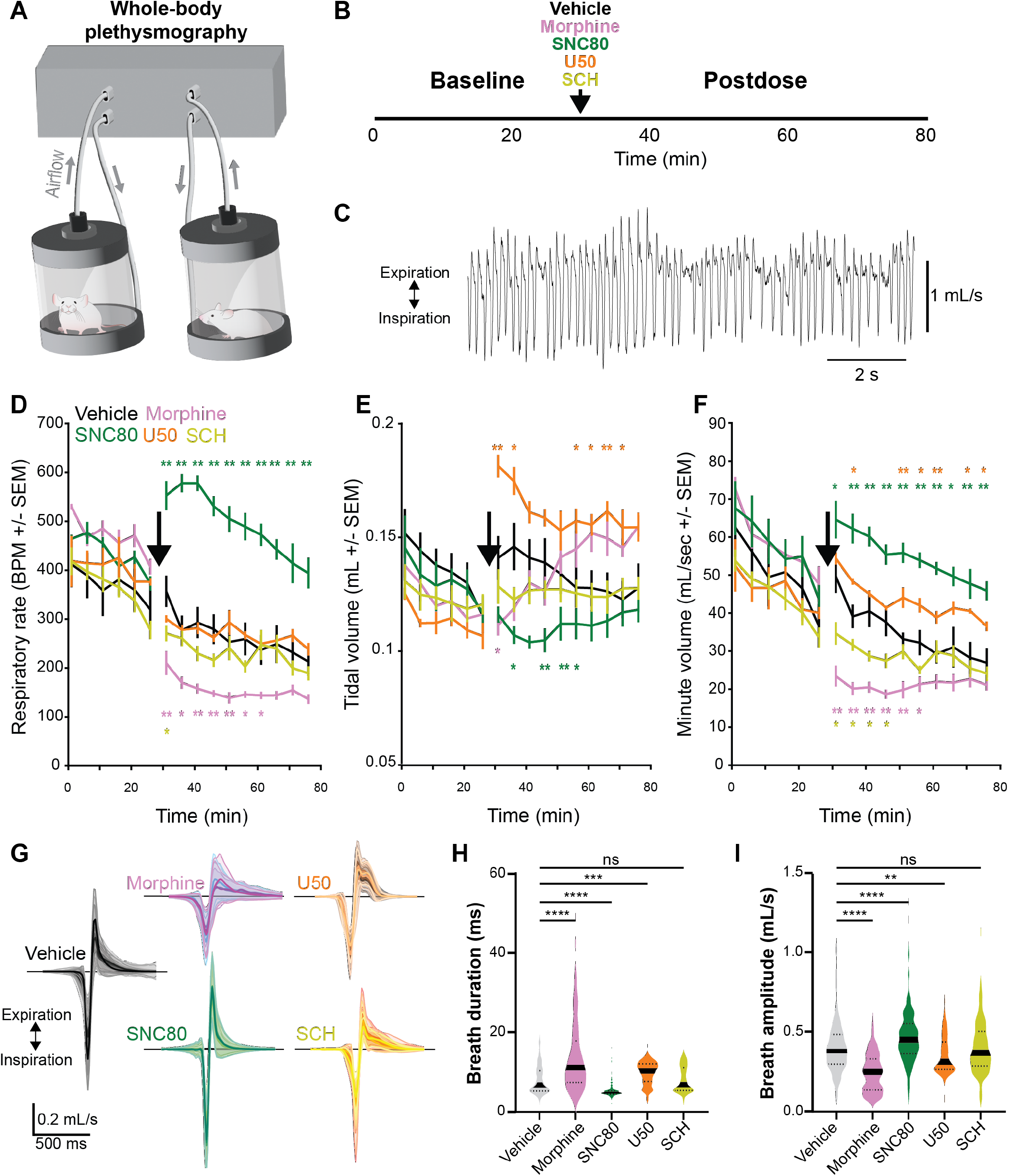
Effect of MOR, DOR, KOR, and NOP agonism on respiration. **(A)** Whole-body plethysmography apparatus enabling respiratory measurements in freely moving rodents. **(B)** Experimental timeline. (**C)** Example respiratory trace during baseline. Respiratory rate (**D)**, tidal volume **(E)**, and minute volume **(F)** of mice during baseline (beginning at minute 0 to the arrow) and postdose (from arrow to the end) of vehicle (black), 20 mg/kg morphine (magenta), 5 mg/kg SNC80 (green), 6 mg/kg (±)-U-50488 (orange), or 10 mg/kg SCN221510 (yellow). Significance is indicated by (*) p < 0.05 and (**) p < 0.01. **(G)** Breath shapes identified during postdose for each drug administered. Breath duration (H) and amplitude (I) measured during the postdose for each drug. (**) indicates p < 0.01, (***) p < 0.001, and (****) p < 0.0001.

KOR and NOP agonists had little to no effect on respiratory rate. However, administration of the KOR agonist correlated with a significant increase in tidal volume, resulting in an overall increase in minute ventilation (**Fig 6E,F**). Close examination of the KOR-induced breath shapes revealed a significant rise in breath duration and decrease in breath amplitude, indicating that although KOR may not impact the respiratory rate directly, KOR activation in respiratory circuits likely alters respiratory patterning (**Fig 6G,H**).

The DOR agonist SNC80, however, elicited a rapid and sustained increase in respiratory rate (**Fig 6D**), decrease in breath duration (**Fig 6G,H**), and increase in inspiratory amplitude (**Fig 6I**). Therefore, DOR and MOR agonists produce opposite effects on respiration, which likely represents the functional consequence of their segregated expression in brainstem sensory-motor circuits.

## Discussion

We found extensive MOR, DOR, KOR, and NOP expression across diverse brainstem nuclei, implicated all four receptors in trigeminal somatosensory-motor (**Figs 2-3**), auditory (**Fig 4**), and respiratory behaviors (**Fig 5-6**). Our quantitative anatomical analysis reveals that all brainstem circuits express at least one OR. Notably, most structures display unique OR expression profiles with few strong correlations or anticorrelations between them (**Fig 1**). Examination of MOR and DOR co-expression showed that, even when present in the same nucleus, these two receptors are predominantly expressed by different neurons (**Figs 2,4**,**5**). Further, we identified function-specific patterns of OR expression and general organizational rules that maintain across brainstem functional modalities. Of note, our behavioral analysis shows that MOR and DOR signaling can cause opposite effects on respiratory circuits, suggesting that MOR and DOR may generally act in opposition. We thus conjecture that endogenous opioid signaling in braintem affects not only nociception, reward, and respiration, but the entire array of animal behaviors.

### Opioid receptor brainstem expression profiles

It is well known that MOR and KOR are densely expressed in key forebrain structures such as MOR in MHb and KOR in Claust; however, brainstem nuclei like IP and SNr show the second-highest expression MOR and KOR, respectively. Further, the greatest overlap of MOR and KOR expression is found in brainstem nuclei including SpVC, SpVM, PaV, LPB, KF, PAG subregions, and VTA.

Secondary sensory areas (e.g., DCN, SpVC, dorsal SpVO, dorsal SpVIr, NTS, and PaV), motoneurons (e.g., NA, X, VII, XII, VTT), and many subregions of the reticular formation, PAG, SC, and others all contain MOR+ neurons. High expression of MOR often coincides with low, but significant, KOR expression, particularly in cranial sensory and motor nuclei (e.g., dorsal SpVO, dorsal SpVIr, XII, VII). Similar to MOR and KOR, MOR and DOR often overlap. However, we rarely find them expressed in the same neurons. A notable exception is the dorsal part of PrV, a critical node in the thalamocortical circuit for jaw somatosensation, where we found that over half of neurons expressing MOR or DOR were MOR+/DOR+. A second notable exception for MOR and DOR coexpression is the superior olive subregions and VTT motoneurons, where many of the DOR+ neurons also express MOR. Future studies assessing the impact of MOR and DOR agonists on auditory reflexes may reveal the functional significance of this coexpression.

Finally, NOP is expressed broadly across the brainstem, with particularly high expression in the deep cerebellar nuclei, IP, LC, and pFRG. Nuclei with high NOP expression levels tend to express at least one other OR, suggesting that NOP may act to co-modulate neurotransmission with MOR, DOR, and/or KOR in the brainstem. In line with this hypothesis, coadministration of NOP antagonists can modulate the effects of MOR agonists such as morphine, including attenuation of morphine tolerance (Toll et al., 2016).

### μ opioid receptor agonists likely alter respiration via multiple brainstem regions

Exogenous opioid administration depresses respiration, and overdose can lead to cardiorespiratory failure and death (Dahan et al., 2010; Mercer Lindsay and Scherrer, 2019). The specific neuroanatomical location underlying OIRD has remained a topic of debate, partly due to the complexity of brainstem respiratory circuits that span many nuclei and regions (Feldman and Del Negro, 2013; Montandon and Horner, 2014; Del Negro et al., 2018). It is widely assumed that, among the four receptors, only MOR contributes to OIRD (Shook et al., 1990; Greer et al., 1995; Lalley, 2008). Crucially, our findings reveal MOR expression throughout important respiratory brainstem circuits, including two of the three CPGs (**Figs 5, S4**).

Given the stereotypical changes in respiratory rate and inspiratory time associated with opioid treatment, OIRD research tends to focus on rhythmogenesis (Shook et al., 1990; Ramirez et al., 2021). We, and others, identified MOR expression in a subset of preBötC neurons, implicating the preBötC as a critical site driving OIRD, especially in light of behavioral studies showing that virally deleting MOR expression from preBötC reduces the severity of OIRD (Montandon et al., 2011; Montandon and Horner, 2014; Bachmutsky et al., 2020; Varga et al., 2020). Notably, the distinct lack of other ORs in this nucleus further reinforces the notion that MOR activation alone drives the preBötC-mediated component of OIRD.

Of the non-CPG nuclei directly linked to respiratory modulation, PB and the closely associated KF stand out as potential sites participating in OIRD (Bachmutsky et al., 2020; Varga et al., 2020). Both regions process respiratory information from the periphery and modulate respiration via projections to CPGs (Varga et al., 2020). KF plays a specific role in rate and pattern stabilization (Varga et al., 2020). As with preBötC, genetic deletion of MOR expression in KF/LPB partially rescues OIRD (Varga et al., 2020; Liu et al., 2021). In both KF and PB, we observed significant MOR expression, as well as higher expression levels of KOR, NOP, and DOR than in preBötC. Crucially, the MOR deletion studies do not demonstrate complete recovery of respiration to baseline levels (Bachmutsky et al., 2020; Varga et al., 2020; Liu et al., 2021). Our findings that MOR is highly expressed in almost all the respiratory nuclei could explain this incomplete recovery. Instead of occurring at one nucleus, OIRD more likely stems from MOR agonism across the respiratory brainstem, forcing this robust system to falter. That much of the respiratory brainstem expresses no ORs other than MOR corroborates previous work investigating the respiratory impact of various OR agonists. In line with our behavioral findings, other studies show that KOR- and NOP-specific agonists minimally affect respiratory parameters, while conflicting studies report that DOR activation increases, decreases, or does not impact respiratory rate (Shook et al., 1990; Dosaka-Akita et al., 1993; Greer et al., 1995; Su et al., 1998; Gallantine and Meert, 2005; Jutkiewicz et al., 2006; Ko et al., 2009; Linz et al., 2014). We find that KOR and DOR agonists affect the breath shape (i.e., inspiratory and expiratory patterns). Based on our data, we propose that coadministration of a DOR agonist could counter MOR-induced respiratory depression; however, future studies would be needed to determine the details given conflicting studies regarding the impact of different DOR agonists on respiration mentioned above. In contrast to DOR, KOR agonism modulated breath shapes similarly to MOR in that breaths were longer and their amplitudes smaller, suggesting that activating KOR in addition to MOR could worsen OIRD.

Given that activation of each ORs resulted in distinct, though subtle, changes to the average breath waveform (**Fig 6**), expression of non-MOR ORs in the respiratory nuclei may affect respiration, though not significantly enough to cause observable changes to the overall pattern of breathing. However, the mechanisms underlying opioid-mediated respiratory modulation requires further investigation. Notably, these effects could also originate in non-brainstem components of the respiratory neural circuitry (i.e., the cortex or airway neural pathways).

Altogether, we have shown that MOR, DOR, KOR, and NOP are prevalent across brainstem circuits, in which these receptors modulate a variety of crucial functions beyond pain and respiration. Given the vast array of behaviors modulated by brainstem circuits, opioids here likely produce an array of effects insufficiently considered, in animal research and potentially in the clinic as well.

## Methods

### Mice

All procedures followed animal care guidelines approved by the Stanford University Administrative Panel on Laboratory Animal Care (APLAC) and University of North Carolina Institutional Animal Care and Use Committee (IACUC), and abided by the recommendations of the International Association for the Study of Pain (IASP). Mice were housed 2-5 per cage in a temperature-controlled environment with a 12 hr light/dark cycle and ad libitum access to food and water. We used male and female mice between 5 and 8 weeks old for all experiments. We used 10 *Oprd1*^GFP/GFP^ mice (Scherrer et al., 2006), 12 *Oprm1*^mCherry/mCherry^ mice (Erbs et al., 2015), 3 *Oprm1*^Cre/Cre^ mice (Bailly et al., 2020), 5 *Oprl1*^YFP-flox/YFP-flox^ mice (Mann et al. 2019), 6 *Oprk1*^tdTomato/tdTomato^ mice (Chen et al., 2020), and 10 C57BL/6 mice.

### Histology and image processing

Mice were deeply anesthetized with pentobarbital and perfused transcardially with 0.1 M phosphate-buffered saline (PBS) followed by 4% formaldehyde solution (Thermo Fisher BP531-500) in 0.1 M phosphate buffer (PB). Brains were dissected, post-fixed for 4 hr at 4°C, and cryoprotected overnight in 30% sucrose in PBS. Frozen brains were then cut at 40 µm using a Leica Cryostat 3050S or a Tanner Scientific TN50 Cryostat.

### Coronal brainstem section processing

Coronal sections were incubated with 5% NDST (0.3% Triton X-100 solution in 0.1 M PBS plus 5% normal donkey serum) for 1 hr. Primary antibodies were diluted in the same solution, and incubated with brain sections overnight at 4°C. After washing the primary antibody 3 times for 10-15 min with 0.3% Triton X-100 solution in 0.1 M PBS, sections were incubated with secondary antibody solution in 1% NDST solution at room temperature for 2 hr. After washing with PB 3 times for 5-10 min, sections were mounted in glass slides using Fluoromount-G (Southern Biotech). Images were acquired with either a confocal (Zeiss 780) or epifluorescence (Zeiss Axio Imager Z1) microscope.

### Sagittal brainstem section processing

The quantification and 3D brainstem atlas were created from 40 µm sagittal sections. After sectioning on a cryostat (Tanner Scientific TN50), sections were immediately mounted onto slides. After drying at room temperature overnight, slides were washed with PB before applying 2% NDST for 30 min. Primary antibodies were diluted in 2% NDST and left on the slides overnight at room temperature. Slides were then washed 2 times with PB and then incubated in secondary antibodies in 2% NDST for 2 hr. Slides were washed once in PB, then a fluorescent Nissl stain was applied (1:200 in 2% NDST, Thermo Fisher N21479 for MORmCh and KORtdTomato or N21483 for DORGFP and NOPYFP) for 20 min. Slides were washed 2 times for 10 min, then coverslipped with Fluoromount (Southern Biotech). Whole-brain slide scanning (10X, 0.25 numerical aperture) was performed on a Zeiss Axio Scan Z1 with a 16-bit AxioCam HRm. Image acquisition was performed using the ZEN software (Zeiss) and stitched with Zen macros. Image acquisition of the opioid receptor channel was calibrated to the entire brain to ensure the same exposure and subsequent image processing for all sections.

### Full brainstem quantification

Quantification was performed using custom MATLAB scripts that compute mean fluorescence of each OR channel based on nucleic boundaries defined by the user from the NeuroTrace image. In order to compare between mouse lines, fluorescence was normalized by setting the I/II&SpVM (effectively lamina I and II of SpVC and the SpVI/SpVC transition zone) at 100% and the background fluorescence a value of 0%. All values are reported relative to the fluorescent range between the I/II&SpVM and background.

Alignment of sections to a 3D volume was performed first in QuickNII (Puchades et al., 2019) using the NeuroTrace images. Fluorescence in the OR channel was detected using ilastik software (Berg et al., 2019). Subsequent alignment to the Allen Common Coordinate Framework (CCF) was performed in Nutil (Yates et al. 2019, Groeneboom et al. 2020). 3D rendering and extraction of pixels per Allen CCF region was performed using BrainMesh (Yaoyao, 2020). Structures that were not aligned in QuickNII due to variability between mice were adjusted using custom boundaries based on NeuroTrace images. Custom MATLAB scripts were used to fully analyze the density and distribution of labeling throughout structures.

### Antibodies

We used the following primary antibodies: anti-GFP, Invitrogen A11122 (rabbit, 1:1000); anti-RFP, Rockland 600-901-379S [chicken, 1:1000 (coronal) or 1:500 (sagittal)]. We used the following secondary antibodies: Jackson Immuno Donkey anti-Chicken 488 (703-545-155); Invitrogen Donkey anti-Rabbit 488 (A21206); Jackson Immuno Donkey anti-Chicken 594 (703-586-155).

### Drug administration

Prior to immunohistochemistry experiments in *Oprd1*^GFP/GFP^ mice, SNC80 (10 mg/kg, Tocris 0764) was subcutaneously injected via a 30 G needle attached to an insulin syringe inserted through the back skin into lightly restrained unanesthetized mice. Behavioral effects of the drug were confirmed 15 min post-injection. The animals were sacrificed 2 hr post-injection.

### Behavior

#### Plethysmography

Respiratory behavior experiments were performed using Buxco Small Animal Whole Body Plethysmography. Adult wild-type mice were placed in the chamber for 30 min to allow the animal to acclimate to the chamber and obtain a baseline respiratory rate. IP injections of saline, 20 mg/kg morphine, 5 mg/kg SNC80, 6 mg/kg (±)-U-50488, or 10 mg/kg SCN221510 were then performed, and the mice were placed in the chamber for another 60 min. Data were exported from the system as voltage traces representing airflow within the chambers and analyzed in MATLAB. To avoid potential artifacts, the first and last 5 min of the post-injection time course were not analyzed.

---

**Table 1.**
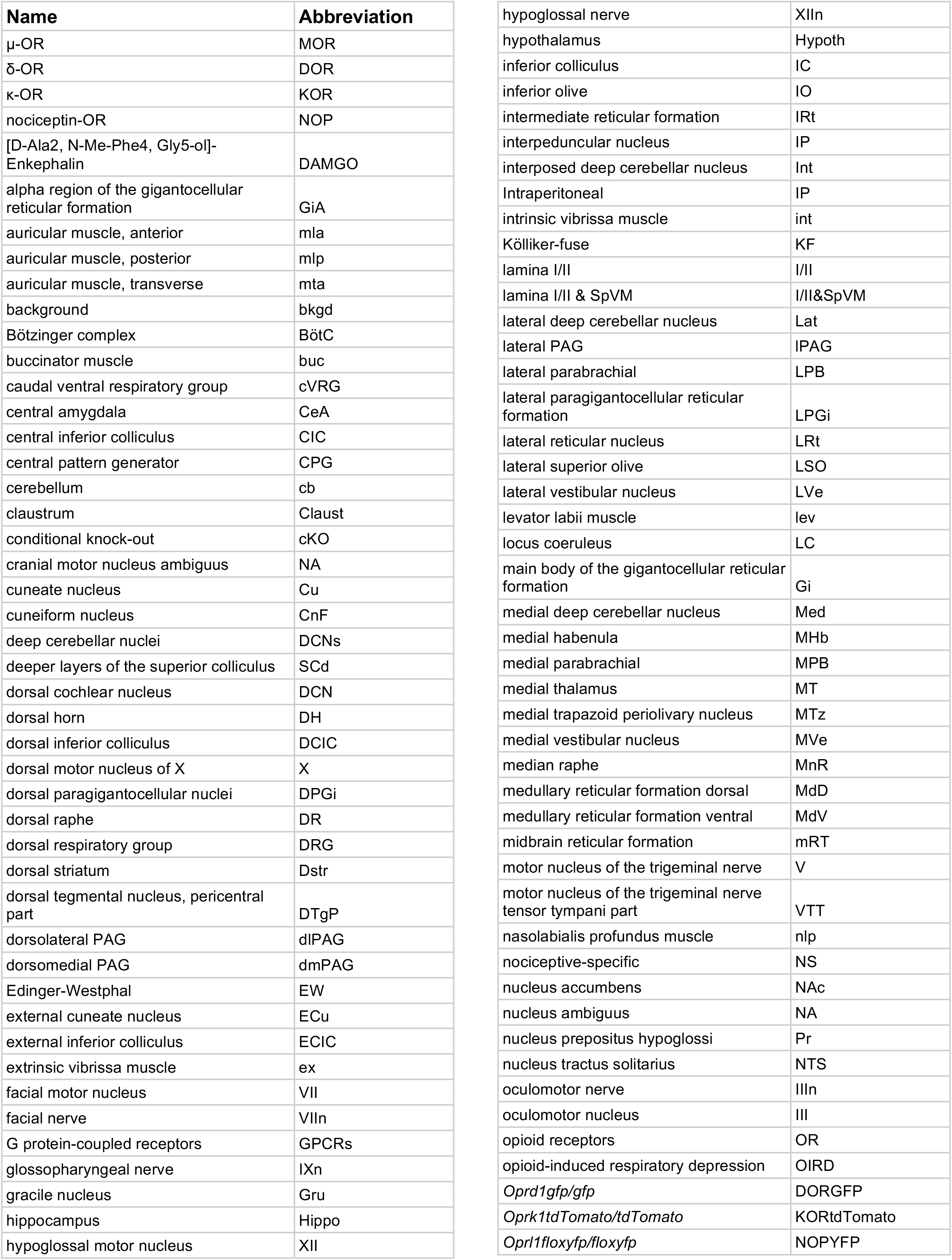
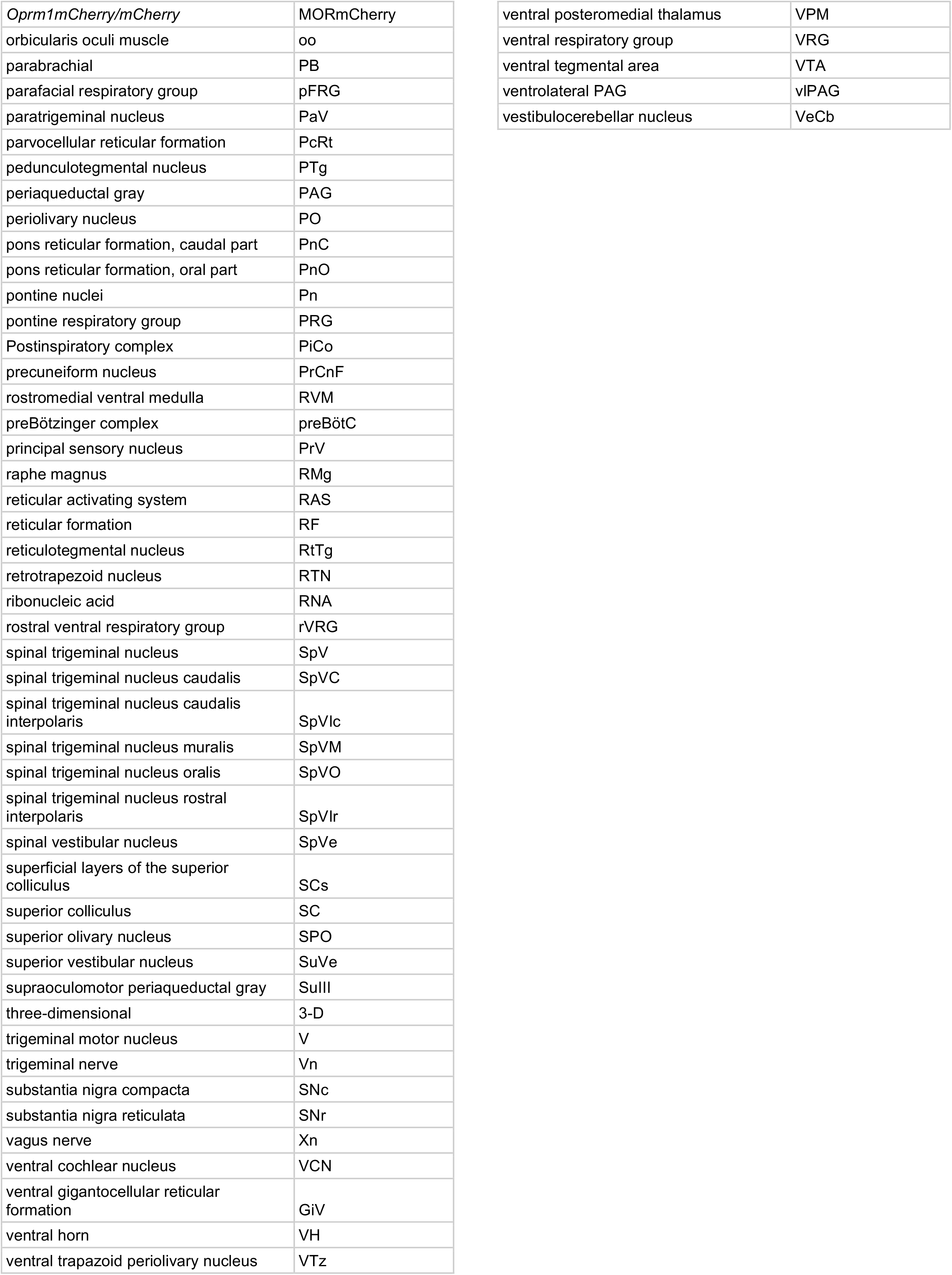
Nomenclature reference.

